# Long-read RNA sequencing reveals widespread sex-specific alternative splicing in threespine stickleback fish

**DOI:** 10.1101/2020.11.12.380428

**Authors:** Alice S. Naftaly, Shana Pau, Michael A. White

## Abstract

Alternate isoforms contribute immensely to phenotypic diversity across eukaryotes. While short read RNA-sequencing has increased our understanding of isoform diversity, it is challenging to accurately detect full-length transcripts, preventing the identification of many alternate isoforms. Long-read sequencing technologies have made it possible to sequence full length alternative transcripts, accurately characterizing alternative splicing events, alternate transcription start and end sites, and differences in UTR regions. Here, we utilize PacBio long read RNA-sequencing (Iso-Seq) to examine the transcriptomes of five tissues in threespine stickleback fish (*Gasterosteus aculeatus*), a widely used genetic model species. The threespine stickleback fish has a refined genome assembly with gene annotations that are based on short-read RNA sequencing and predictions from coding sequence of other species. This suggests some of the existing annotations may be inaccurate or alternative transcripts may not be fully characterized. Using Iso-Seq we detected thousands of novel isoforms, indicating many isoforms are absent in the current Ensembl gene annotations. In addition, we refined many of the existing annotations within the genome. We noted many improperly positioned transcription start sites that were refined with long-read sequencing. The Iso-Seq predicted transcription start sites were more accurate, verified through ATAC-seq. We were also able to detect many alternative splicing events between sexes and across tissues. We found a substantial number of genes in both somatic and gonad tissue that had sex-specific isoforms. Our study highlights the power of long-read sequencing to study the complexity of transcriptomes, greatly improving genomic resources for the threespine stickleback fish.

## Introduction

The ability to generate alternative isoforms from a finite number of genes is a widespread phenomenon across eukaryotes that has been hypothesized to play a key role in the evolution of phenotypic diversity (reviewed in Graveley 2001; Keren et al. 2010; Baralle and Giudice 2017). Alternative isoforms can arise through multiple mechanisms. One mechanism is to alter the coding sequence of the transcript through alternative splicing. This can be achieved through the retention of introns, the inclusion or exclusion of entire exons, or through the usage of alternative splice sites within exons (reviewed in Smith et al. 1989; Keren et al. 2010). Isoform diversity can also be increased through the inclusion of alternate transcription start sites (TSSs) or transcription termination sites (TTSs), leading to differences in the 5’ or 3’ untranslated regions (UTRs). These variants do not alter the underlying coding sequence, but can alter transcriptional regulation and underlying stability of the mRNA transcript (Gupta et al. 2014; Wang et al. 2016b; Zhang et al. 2017).

Short read RNA-seq has greatly expanded our ability to survey the complexity of transcriptomes (reviewed in Costa et al. 2010), including the computational prediction of alternative splicing events (Trapnell et al. 2009; Anders et al. 2012; Kim et al. 2015; Kim et al. 2019). However, RNA-seq cannot accurately detect all isoforms present in a transcriptome. In order to detect all splice junctions among alternative isoforms there must be sufficient read depth at alternative exon-exon boundaries (Bryant et al. 2012; Steijger et al. 2013). For experiments with insufficient read coverage, lowly expressed isoforms are challenging to recover and predict (reviewed in Conesa et al. 2016). Alternative isoform identification is further confounded by short reads mapping to multiple isoforms which can lead to collapse into a single isoform. To properly differentiate alternative transcripts, full-length sequencing of transcripts is necessary (Steijger et al. 2013; Wang et al. 2016a).

Long-read sequencing technologies have made it possible to sequence a single full-length transcript with high accuracy (Wang et al. 2016a; Wang et al. 2019).With sufficient sequencing coverage, all isoforms can be identified unambiguously, classifying the complete catalog of splice junctions and alternate TSSs and TTSs (Wang et al. 2019). This technology has been successfully applied to multiple species of plants and animals (Sharon et al. 2013; Abdel-Ghany et al. 2016; Wang et al. 2016a; Cheng et al. 2017; Kuo et al. 2017; Li et al. 2018; Nudelman et al. 2018; Deslattes Mays et al. 2019; Zhang et al. 2019). In each case, long-read RNA sequencing has helped refine existing gene annotations as well as characterized pervasive alternative splicing among tissues (Abdel-Ghany et al. 2016; Wang et al. 2016a; Kuo et al. 2017; Li et al. 2018; Zhang et al. 2019).

Although transcriptome complexity has been increasingly studied at the tissue level, comparatively little is known about the sex-specificity of isoforms. Sexual dimorphism in alternative splicing may be important in regulating many of the phenotypic differences observed between sexes. For instance, male and female somatic differentiation is controlled by alternatively spliced transcripts of the *doublesex* gene (Burtis and Baker 1989). In addition, alternative splicing can be a mechanism to resolve intralocus sexual antagonism, where the expression of a gene is beneficial to one sex yet harmful to the other (reviewed in Ellegren and Parsch 2007; Stewart et al. 2010). Alternative splicing could allow antagonistic exons to be restricted to a single sex or alternative TSSs and TTSs could create opportunities for sex-specific transcriptional regulation. At a genome level there is growing evidence that alternative splicing is widespread between sexes (McIntyre et al. 2006; Blekhman et al. 2010; Brown et al. 2014; Gibilisco et al. 2016; Rogers et al. 2020). However, all surveys have utilized either short-read RNA-seq or microarray probes targeting known transcripts. This raises the possibility that the true amount of alternative splicing between sexes may be underestimated.

Here we use PacBio Iso-Seq long-read sequencing of transcriptomes isolated from threespine stickleback fish (*Gasterosteus aculeatus*) to explore the extent of alternative isoforms across tissues and between sexes. Threespine stickleback fish are an emerging genetic model system for evolutionary biology, ecology, behavior, physiology, and toxicology (Bell and Foster 1994; Kingsley 2003; Barber and Nettlseship 2010; Hendry et al. 2013). There is a high-quality genome sequence that has undergone several rounds of revision (Peichel et al. 2001; Jones et al. 2012a; Glazer et al. 2015; Peichel et al. 2017). Although the genome sequence has been curated well, the gene annotations are based entirely from the Ensembl annotation pipeline (Yates et al. 2020), incorporating short expressed sequencing tags (ESTs), publicly available short-read RNA sequences, and known homology from other organisms (Yates et al. 2020). In an effort to thoroughly survey the extent of alternative isoforms, we sequenced the transcriptomes of five tissues to high coverage in both sexes (liver, brain, pronephros, testis, and ovary). Using Iso-Seq, we were able to refine many of the Ensembl annotations. Our survey also detected thousands of novel isoforms compared to the Ensembl annotations. Between sexes, a large percentage of genes had male or female specific isoforms, highlighting the overall power of long-read RNA sequencing to characterize transcriptional diversity. These transcriptomes will be an important resource in exploring the overall contribution of alternative splicing to sexual dimorphism as threespine stickleback fish exhibit pronounced phenotypic differences between males and females (Kitano et al. 2007; Leinonen et al. 2011; Kotrschal et al. 2012; McGee and Wainwright 2013).

## Results

### PacBio long-read sequencing identified several thousand isoforms

On average, 618,000 CCS reads per tissue were produced from the raw subreads (Supplemental Tables 1 and 2). After filtering, full-length classification, and polishing, each tissue had an average of 41,000 high quality reads (Supplemental Table 1). Using SQANTI filtering (see methods), we recovered a final Iso-Seq transcriptome composed of 26,432 isoforms (13,703 genes; annotated genes: 7,754; novel genes: 5,949; Supplemental Files 1-4, Supplemental Table 3). In comparison, the Ensembl transcriptome (build 97) had 22,443 genes and 29,245 isoforms. Although we found a smaller number of genes in the Iso-seq transcriptome, we identified a similar number of total isoforms suggesting that alternative splicing and/or alternative TSSs and TTSs may be more pervasive than predicted through the Ensembl annotations. Consistent with this, we identified 18,271 novel isoforms that did not match any of the previously annotated Ensembl isoforms. Furthermore, within single genes we also observed a greater breadth of alternative isoforms. There was a greater proportion of genes that had two or more isoforms in the Iso-Seq transcriptome compared to the Ensembl transcriptome (Ensembl: 24%; Iso-Seq: 31%; Supplemental Figure 1).

We used short-read RNA sequencing to verify the accuracy of alternative splicing among the Iso-seq isoforms. We sequenced each tissue to high coverage in order to target a complete set of alternative splice junctions (~166 million reads per tissue were produced; Supplemental Table 4). We searched for the presence of uniquely mapping and multimapping short reads that spanned the splice junctions of isoforms with two or more exons (17,853 isoforms). The short-read sequencing revealed the Iso-seq sequencing was highly accurate at detecting alternative isoforms. A majority of the isoforms (16,826 isoforms, 94% of isoforms with two or more exons) had all splice junctions confirmed by short read sequencing through both uniquely mapping reads and multimapping reads. Only 321 isoforms (2%) had no short reads supporting the splice junctions. The remaining 706 isoforms (4%) had one or more splice junctions not confirmed by the short-read data.

We assessed transcriptome completeness using benchmarking universal single-copy orthologs (BUSCO). BUSCO utilizes essential single copy orthologs that are expected to be present in complete transcriptomes (Simao et al. 2015; Seppey et al. 2019). We compared the existing threespine stickleback Ensembl transcriptome and the new Iso-Seq transcriptomes to two databases, metazoa with 978 genes and actinopterygii with 4,584 genes. The Ensembl transcriptome contained mostly complete orthologs. 97% were complete orthologs in metazoa (single copy orthologs: 661; duplicated orthologs: 285; Supplemental Figure 2) and 98% were complete orthologs in actinopterygii (single copy orthologs: 3,416; duplicated orthologs: 1,053; Supplemental Figure 2). The Iso-Seq transcriptome had fewer complete orthologs. 61% of the metazoan gene sets were complete (single copy orthologs: 255; duplicated orthologs: 341; Supplemental Figure 2) and 35% of actinopterygian gene sets were complete (single copy orthologs: 3,416; duplicated orthologs: 1,053; Supplemental Figure 2).

One possibility why we did not capture more single copy orthologs was that our libraries may not have been sequenced to an adequate depth. To test this, we used a subsampling approach (Workman et al. 2018). We subsampled the CCS reads of each individual tissue and compared those reads to all of the predicted isoforms for that tissue. We found we were able to recover at least 90% of the predicted isoforms with only 35% to 85% of the original CCS reads (Supplemental Figure 3). This indicates that each library was saturated with reads and including additional sequencing would not greatly increase the transcriptome completeness. Therefore, the lack of completeness in single copy orthologs in our dataset likely reflects the limited number of tissues we sequenced.

### Full-length isoform sequencing refined many of the previously predicted transcription start and end sites

SQANTI characterizes isoforms into nine different categories based on the splice junctions between exons (Tardaguila et al. 2018). We condensed these nine categories into three broad classes: Ensembl isoform matches, novel isoforms, and novel genes. Of the 26,432 total protein coding isoforms, only 6,410 exactly matched the Ensembl predicted splicing (25%; Figure 1, Table 1). The remainder of the isoforms that overlapped Ensembl annotations did not match the existing splicing annotations fully and likely represent alternative splicing events that exclude some internal exons (1,551 isoforms; Figure 1, Table 1). Interestingly, we found that a majority of isoforms that fully matched internal Ensembl splice junctions had differences in TSS’s and TTS’s (full splice match isoforms that did not match annotated TSS: 99%; full splice match isoforms that did not match annotated TTS: 98%). On average, the full splice match isoforms had TSS’s that were 99 bp further upstream of the annotated TSS (Figure 2). This pattern was more pronounced in the TTSs where the average distance from the Ensembl TTS was 500 bp upstream (Figure 2).

**Figure 1.**
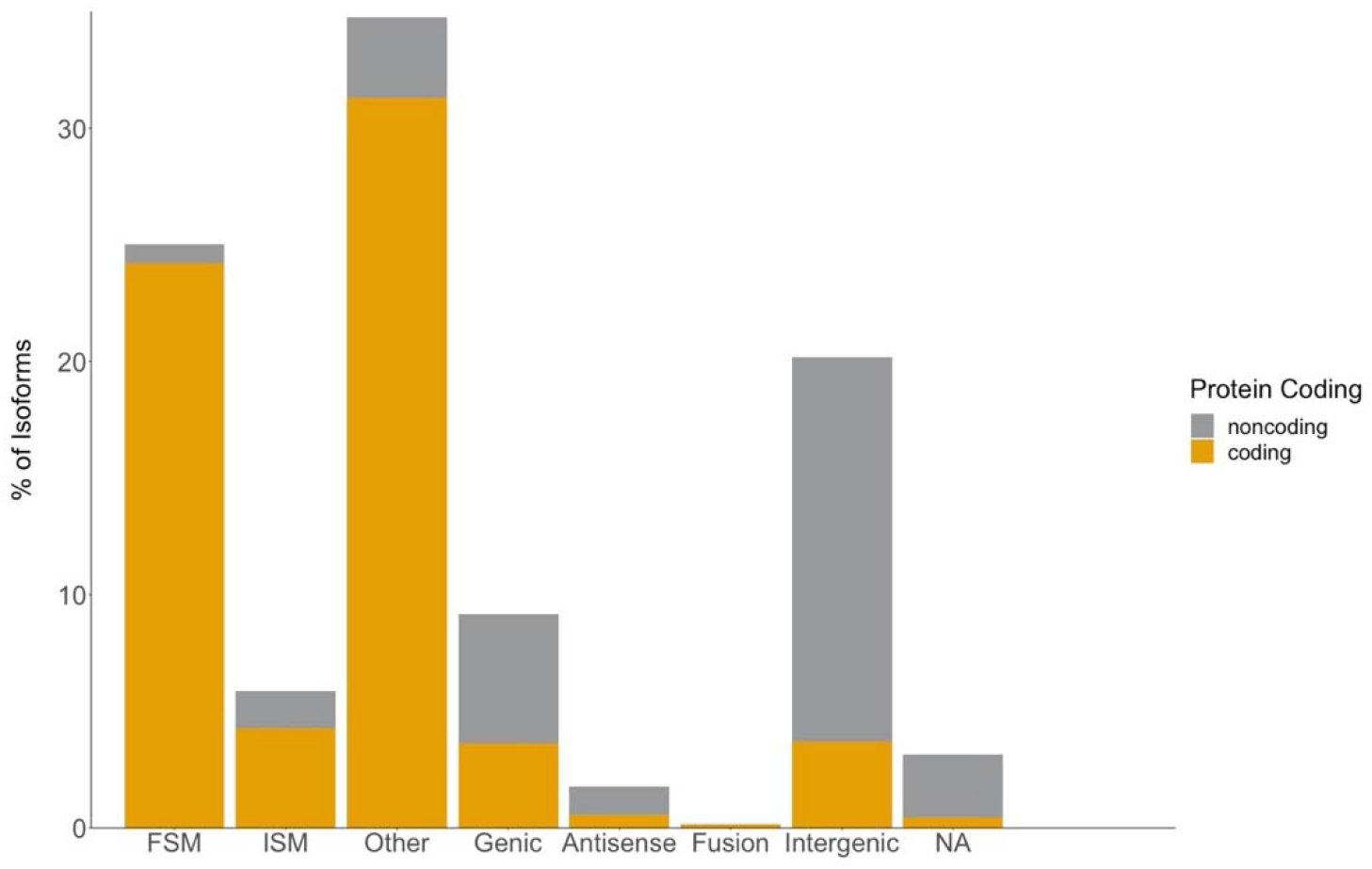
In-depth characterization of isoforms by SQANTI for Iso-Seq transcriptome. Isoforms are classified as one of nine different splice categories by SQANTI: FSM (full splice matches), ISM (incomplete splice match), NIC (novel in catalog), NNC (novel not in catalog), Genic Genomic, Antisense, Fusion, Intergenic, and Genic Intron. The isoforms in each splice category are divided into predicted protein coding isoforms (gold) and predicted non-protein coding isoforms (grey).

**Table 1:** Characterization of Iso-Seq transcripts.

| **Isoform Category** | **Isoform Count** |
| --- | --- |
| Isoform match with Ensembl: protein coding |  |
| Full splice match | 6,401 |
| Incomplete splice match | 1,125 |
| Isoform match with Ensembl: non-protein coding |  |
| Full splice match | 209 |
| Incomplete splice match | 426 |
| Novel Isoform: protein coding |  |
| Genic | 964 |
| Fusion | 39 |
| Other | 8,273 |
| Novel Isoform: non-protein coding |  |
| Genic | 1,461 |
| Fusion | 2 |
| Other | 907 |
| Novel Gene: protein coding |  |
| Intergenic | 981 |
| Antisense | 152 |
| Intronic | 123 |
| Novel Gene: non-protein coding |  |
| Intergenic | 4,349 |
| Antisense | 316 |
| Intronic | 704 |
| Total Isoforms* | 26,432 |

**Figure 2.**
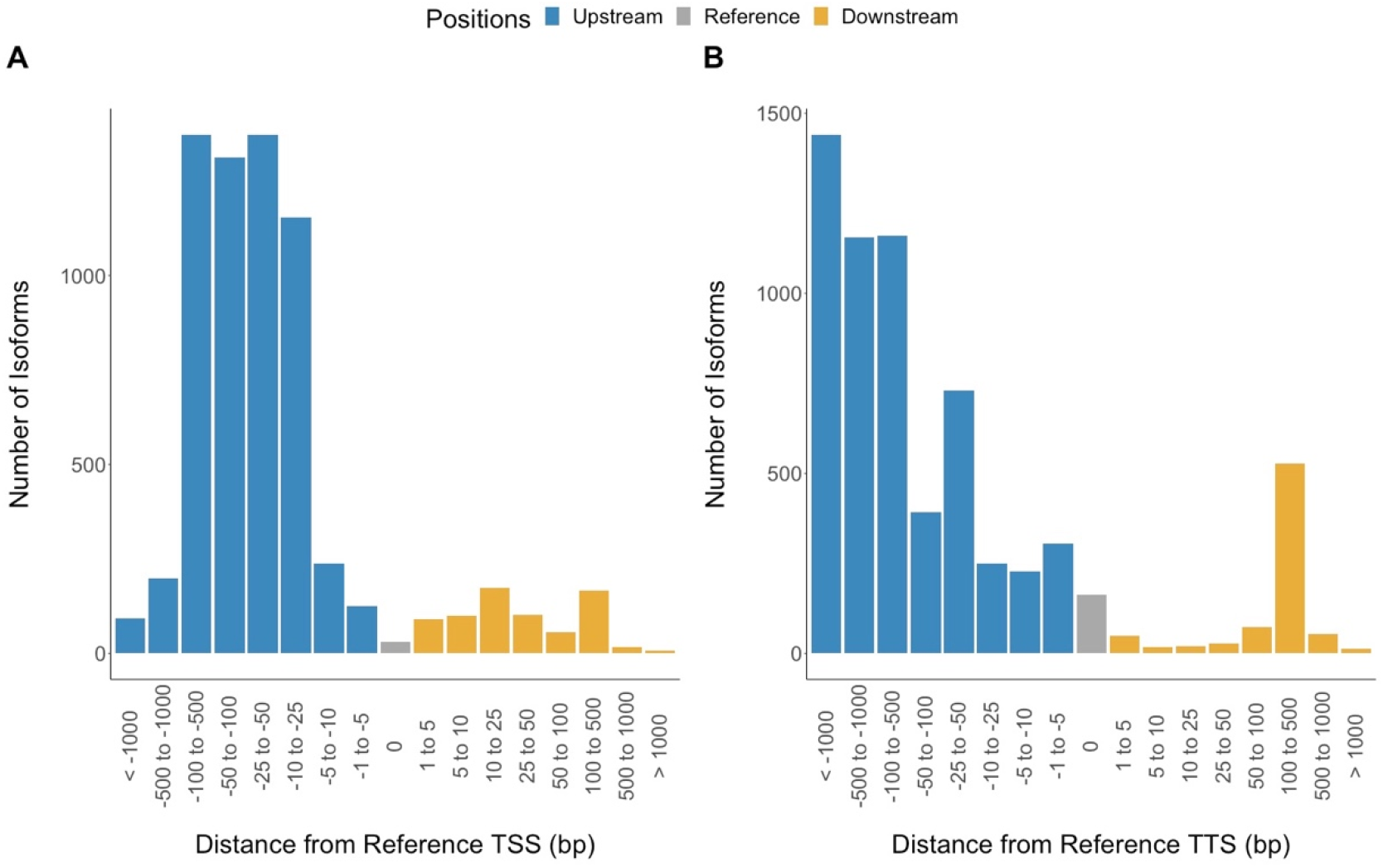
Iso-Seq full splice matches have different transcription start sites (TSS) and transcription termination sites (TTS) compared to Ensembl annotations. Full splice matches are isoforms that have the same exon boundaries as Ensembl transcripts (6,610 isoforms). (A) On average, full splice match isoform TSSs are located 99 bp upstream of the annotated TSS. (B) Full splice match isoform TTSs are located on average 500 bp upstream of the annotated TTS.

We confirmed the shift in TSSs by utilizing ATAC-seq chromatin accessibility data from liver samples. Sequencing reads from ATAC-seq are expected to be enriched around the nucleosome-free region of TSSs (Mavrich et al. 2008; Buenrostro et al. 2013; Meers et al. 2018). We compared the read coverage in the 4 kb surrounding Ensembl annotated TSSs and in the 4 kb surrounding the newly annotated TSSs from Iso-Seq. We found that there is a much higher enrichment of ATAC-seq reads that is narrowed exactly at the TSS identified by Iso-Seq (Figure 3, Supplemental Figure 4). This enrichment was much weaker around the TSS in the Ensembl annotations, indicating that many of the existing annotated TSSs are incorrect and can be refined using full-length isoform sequencing.

**Figure 3.**
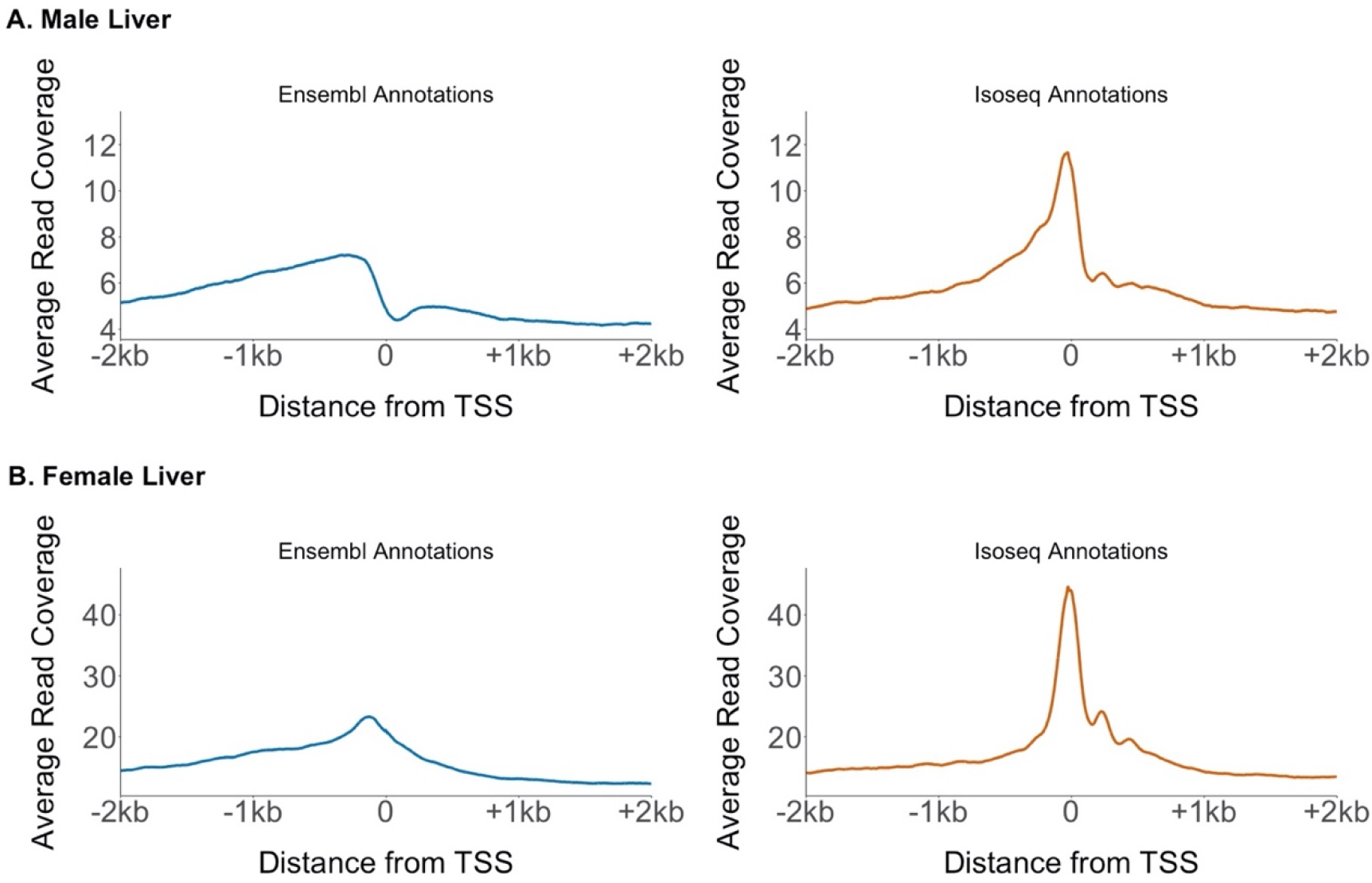
Accessible chromatin is localized in narrow peaks around the Iso-Seq transcription start sites. We compared ATAC-seq read coverage at all Ensembl TSSs and Iso-Seq TSSs across the autosomes. ATAC-seq reads show an enrichment at the Iso-Seq TSS compared to the Ensembl TSS. This indicates a more accurate positioning of the TSS using Iso-Seq. A second male and female replicate is shown in Supplemental Figure 4.

### A majority of the Iso-Seq transcriptome was previously unannotated

Nearly 70% of isoforms detected in the Iso-Seq transcriptome are novel (18,271 isoforms) and fall into the novel isoform or novel gene classes mentioned above. Novel isoforms represent a new isoform of a previously annotated gene in the Ensembl transcriptome where novel genes represent a completely novel gene not previously annotated. A majority of the novel Iso-Seq transcriptome (64%; 11,646) overlaps currently annotated genes, indicating long-read sequencing is able to capture alternative splicing events not readily annotated by current pipelines or is able to identify errors in current exon/intron boundaries (Table 1). The remaining isoforms represented completely novel genes (36%; 6,625), either located entirely within an intergenic region as defined by Ensembl, located entirely within an Ensembl annotated intron, or located on the antisense strand (Table 1). To examine the functions of the novel protein coding isoforms (11,646), we searched for gene ontology matches and similarities to known protein domains. We found 10,107 novel isoforms that had shared sequence identity with at least one known protein domain (Supplemental File 5). Within these results, we looked for GO terms enriched in the novel isoforms. Many of the GO terms were enriched for general components of the cell (e.g. membrane components or organelle components) and also enriched for general molecular functions such as catalytic activity and biogenesis (Supplemental Figure 5, Supplemental Table 5). There was also enrichment in various biological processes such as biological regulation, response to stimulus, reproduction, and immune processes (Supplemental Figure 5, Supplemental Table 5).

### Non-protein coding isoforms are enriched for long non-coding RNAs

We identified a total of 8,374 isoforms that were non-protein coding (32% of the 26,432 total isoforms). This fraction was substantially larger than the proportion of non-protein coding isoforms currently annotated in the Ensembl transcriptome (10%; 2,767 non-protein coding isoforms of 29,245 total isoforms), indicating long-read sequencing may have captured a broader sampling of regulatory non-coding RNAs (ncRNAs). We first examined whether any of our isoforms overlapped with currently annotated regulatory RNAs in the Ensembl transcriptome. We recovered 29% of the previously annotated Ensembl ncRNAs in the Iso-Seq transriptome. Of the 2,767 annotated Ensembl ncRNAs, we found only 209 exactly matched the Iso-Seq annotations (from the Ensembl isoform match: non-protein coding category; Table 1) and 582 partially overlapped an existing ncRNA annotation (distributed among the Ensembl isoform match: non-protein coding category and the novel isoform: non-protein coding category; Table 1).

We characterized the remaining novel non-protein coding isoforms and genes (7,583 total distributed among the novel gene: non-protein coding and novel isoform: non-protein coding categories; Table 1) based on overall length and genome location. Short ncRNAs are typically under 200 bp in length and include miRNAs, endo-siRNAs, and piRNAs (reviewed in Farazi et al. 2008; Pauli et al. 2011). Long ncRNAs (lncRNAs) are greater than 200 bp in length (Mercer et al. 2009). A majority of the novel short and long ncRNAs we identified were classified as intergenic (short ncRNAs: 306 isoforms, 69%; long ncRNAs: 4,135 isoforms, 58%). Far fewer ncRNAs were intronic (short ncRNAs: 44 isoforms, 10%; long ncRNAs: 584 isoforms, 8%) or on the antisense strand (short ncRNAs: 27 isoforms; 6 %; long ncRNAs: 498, 7%). There were 69 (16%) short ncRNAs and 1,920 (27%) long ncRNAs that did not fall into these three regions. These isoforms partially overlapped an annotated isoform on the same strand, likely overlapping an annotated exon. The annotated isoform overlapped was not one of the non-protein coding isoforms annotated in Ensembl.

### Many isoforms have sex-specific alternative splicing

Alternative splicing plays a key role in increasing protein diversity using a limited number of genes. Different isoforms of the same gene can regulate specific developmental and physiological processes (Graveley 2001; Baralle and Giudice 2017). Although alternative splicing has been shown to be important for sex determination in *Drosophila* (Burtis and Baker 1989) and for the development of sex-specific tissues in other species (Telonis-Scott et al. 2009; Gibilisco et al. 2016; Planells et al. 2019), the overall importance of sex-specific alternative splicing remains largely underexplored (McIntyre et al. 2006). This is because detecting full-length isoforms is difficult when using short read RNA-seq. We searched for evidence of sex-specific alternative splicing among the somatic and gonad tissues in our long-read sequencing. We first combined all female tissues (female transcriptome) and all male tissues (male transcriptome) and identified the transcriptomes for each sex (Supplemental Table 3). We found a substantial number of isoforms that were specific to only one sex. Of genes that were expressed in both sexes (4,842 total genes), we found 1,590 (33%) had female-specific isoforms and 2,103 (43%) had male-specific isoforms (Supplemental File 6). Across all genes there were a total of 2,363 female-specific isoforms and 3,664 male-specific isoforms. We characterized whether the alternative isoforms exhibited alternative splicing or whether there were differences in the TSS or TTS that may result in altered transcriptional regulation between the sexes. Nearly half of the isoforms exhibited alternative splicing only in both sexes (female specific: 1,146 isoforms, 49%; male specific: 1,531 isoforms, 42%). The remainder of isoforms had an alternate TSS or TTS (female specific: 425 isoforms, 18%; male specific: 968 isoforms, 26%) or exhibited both alternative splicing and alternate TSS’s/TTS’s (female specific: 792 isoforms, 34%; male specific: 1,165 isoforms, 32%).

We explored whether the alternative isoforms we identified were driven by the inclusion of sex-specific gonad tissues by analyzing the male and female somatic transcriptomes separately (brain, liver, and pronephros combined, Supplemental Table 3). After removal of the gonads, we recovered a similar number of alternative isoforms in females (2,218, 94% of the total sex-specific isoforms recovered from all tissues combined), but a reduced number from males (2,579, 70% of the total sex-specific isoforms recovered from all tissues combined). This suggests that the ovary transcriptome does not contribute greatly to sex-specific alternative isoforms. The testis transcriptome, on the other hand, contains many genes with isoforms unique to males, suggesting this tissue has a much greater transcriptional complexity.

### Pronephros and testis tissue exhibited the greatest degree of alternative splicing

While the majority of genes in all tissues had only one isoform per gene, two tissues had a large proportion of genes that had multiple isoforms per gene (Figure 4A). The male pronephros, female pronephros, and the testis had more than 25% of all genes having two or more isoforms per gene. For the remaining tissues, only 5-15% of all genes had more than two isoforms. Excluding the pronephros, over half of the isoforms identified in the other tissues were annotated in the Ensembl transcriptome (Figure 4B). The pronephros tissues from both sexes had the largest proportion of novel isoforms (female pronephros: 57%; male pronephros: 55%; Figure 4B). Most of the isoforms in each tissue were predicted to be protein coding isoforms (Supplemental Figure 6). The brain had the highest proportion of non-protein coding isoforms (female liver: 19%; male liver: 31%; female brain: 47%; male brain: 37%; female pronephros: 22%; male pronephros: 18%; ovary: 13%; testis: 7%, Supplemental Figure 6).

**Figure 4.**
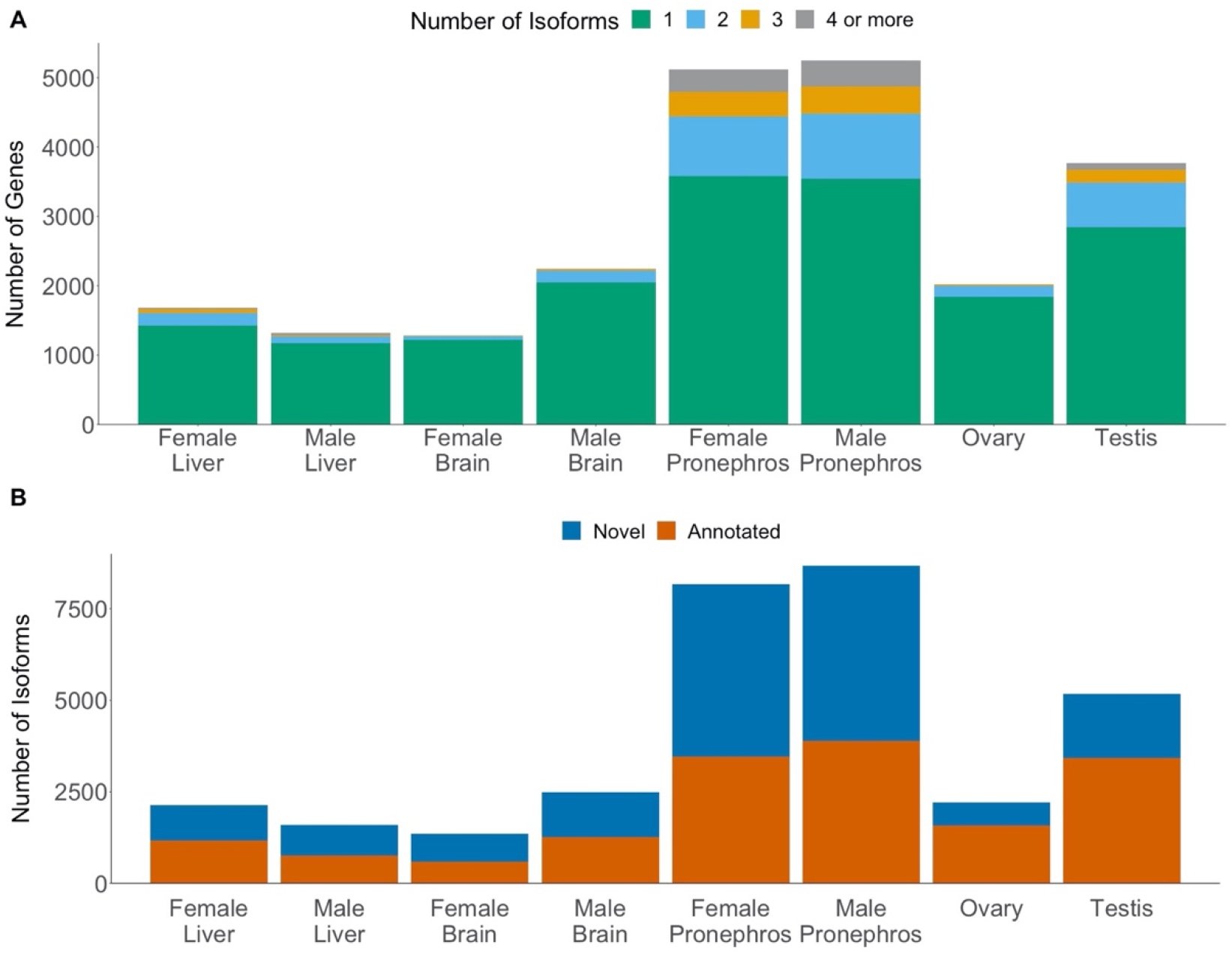
Novel isoforms are found across all tissues and sexes. (A) More than 25% of genes in the testis and pronephros had more than one isoform. For the remaining tissues, less than 15% of the genes had more than one isoform. (B) Over half of the isoforms identified in the pronephros samples are novel isoforms. The testis has the next largest count of novel isoforms.

If a tissue exhibits a disproportionate amount of tissue-specific gene expression, this will raise the number of unique isoforms we detected. To determine if a tissue produces more alternative transcripts relative to the other tissues, we limited the comparison to genes that had shared expression across multiple tissues. Among these genes, both the testis and ovary had the largest number of alternative transcripts (testis: 787 isoforms, 47%; ovary: 146 isoforms, 28%; Supplemental Table 6, Supplemental File 7). A majority of these transcripts were alternatively spliced (brain: 74%; liver: 68%; pronephros: 66%; testis: 44%; ovary: 53%; Supplemental Table 7). The testis had the largest proportion of transcripts that exhibited TSSs and TTSs or both alternative splicing and alternate TSSs and TTSs (Supplemental Table 6). These results suggest that while the pronephros has the largest proportion of tissue specific isoforms, this was largely driven by gene expression, and the testis produced more alternative transcripts relative to other tissues.

## Discussion

### PacBio Iso-Seq greatly improved the gene annotations in threespine stickleback

Using long-read sequencing of several different tissue transcriptomes, we were able to refine the existing isoform annotations across the threespine stickleback genome, adding previously undocumented isoforms, modifying existing splice junctions of isoforms, and correcting previous estimates of TSSs and TESs. The modified splice junctions were highly accurate, verified through deep RNA-seq. We were unable to find reads that covered all splice junctions, but this is not unexpected given short read sequencing often misses or collapses alternative splicing events (Conesa et al. 2016; Wang et al. 2019). Shifts in TSSs also matched patterns of accessible chromatin from ATAC-seq. Correct TSSs will be particularly important for future work in threespine stickleback fish in understanding gene regulation. We were also able to detect many new ncRNAs, similar to patterns seen in other systems using long read technologies (Kuo et al. 2017). These ncRNAs were found across all tissues, however the brain contained the highest proportion. ncRNAs are known to perform a variety of functions in the cell, including housekeeping functions and regulatory functions, and contain ribosomal RNA and transfer RNAs (reviewed in Jacquier 2009; Pauli et al. 2011). Long ncRNAs have been previously reported to be important for the evolution of the human brain and have been associated with specific regions of the brain in mice (Mercer et al. 2008). We also found lncRNAs were prevalent in pronephros tissue. The teleost fish pronephros is an integral component of immune response, containing cytokine producing lymphoid cells (reviewed in Geven and Klaren 2017). In mammals, lncRNAs are important in the development of immune cell lineages (Atianand et al. 2017; reviewed in Ahmad et al. 2020). Our results highlight lncRNAs may have analogous roles in immune development in teleost fish. Additional work will be necessary to functionally validate these transcripts.

Although we were able to capture over 50% of complete metazoan BUSCO orthologs, Iso-Seq from brain, liver, pronephros, and gonads did not approach the total number complete orthologs within the Ensembl transcriptome. This result is not surprising as we only examined five tissues at a single time point, whereas the Ensembl transcriptome compiles data across a wider representation of tissues and also incorporates homology searches to form gene predictions based on coding sequences in other species. Similar patterns of reduced completeness have been reported in other systems where only a few tissues were examined (Workman et al. 2018; Minio et al. 2019). We demonstrated that increasing sequencing depth at an individual tissue would not increase the total number of genes and isoforms detected in our dataset. Therefore, in order to survey transcriptome diversity at a greater number of genes, future work focused on long-read sequencing of additional tissues will be necessary.

### Sex-specific alternative transcripts are ubiquitous across tissues in threespine stickleback

We found that over 30% of alternative transcripts annotated in the Isoseq transcriptome were present in only males or females, regardless of tissue type, suggesting that sex-specific alternative transcripts may be a more widespread phenomenon than previously reported. Sex-specific alternative transcripts have been documented in *Drosophila*, albeit among a smaller proportion of genes than we observed in threespine stickleback fish (McIntyre et al. 2006; Telonis-Scott et al. 2009; Chang et al. 2011; Gibilisco et al. 2016). These surveys utilized exon-specific microarrays or short-read RNA sequencing, raising the possibility that the degree of alternative splicing was underestimated due to limitations in the sequencing technologies. Alternatively, alternative transcripts may be more widespread among genes in threespine stickleback fish, including those that are processed in a sex-specific manner. Indeed, some surveys have suggested that there is a greater number of genes with alternative transcripts in vertebrates compared to invertebrates (Kim et al. 2007). Additional long-read sequencing of transcriptomes will help clarify how extensive alternative transcripts are among taxa.

### Overall tissue complexity varied among threespine stickleback tissues

Transcriptome complexity is often defined as the number of genes present in a given tissue or by the diversity in the transcripts present (Ramskold et al. 2009). In mammals, the brain has been shown to be one of the most transcriptionally complex tissues, with the largest number of isoforms and tissue-specific alternative splicing events (Xu et al. 2002; Kan et al. 2005; Mele et al. 2015). Unlike mammals, we found the threespine stickleback brain has relatively low tissue complexity compared to the other tissues examined. This is likely due to the major differences in brain structure between teleosts and mammals (reviewed in Northcutt 2002).

Instead of high levels of tissue complexity in the brain, we found that the pronephros had the most isoforms followed by the testis. Interestingly, the testis had a much higher proportion of alternatively spliced tissue-specific isoforms than the pronephros. This high level of transcriptome complexity in the testis has also been reported across mammals (Xu et al. 2002; Kan et al. 2005; Ramskold et al. 2009; Schmid et al. 2013; Soumillon et al. 2013). This pattern may be explained by cells undergoing spermatogenesis. Threespine stickleback fish undergo synchronous spermatogenesis, where testes are enriched for cells in the same stage (Craig-Bennett 1931; Borg and Van Veen 1982). The juvenile testes we sequenced contained cells actively undergoing meiosis. In mice, alternative splicing has an important role in meiosis, affeting meiotic progression (Schmid et al. 2013). Key proteins involved in early meiosis also have multiple isoforms (Bellani et al. 2010; Kauppi et al. 2011). Our work offers an interesting parallel to mammalian meiosis and suggests alternative splicing may have a more universal role in spermatogenesis.

The high transcriptome complexity seen in the pronephros is intriguing as the pronephros is present across vertebrates, but only persists into adulthood in amphibians and fish (Smyth et al. 2017). Therefore, there is very little known about this tissue. In fish, the nephritic tissue degenerates over time and the pronephros functions as part of the immune system (reviewed in Geven and Klaren 2017). In mammals, transcriptome complexity of the immune system is high (reviewed in Schaub and Glasmacher 2017), but often is below levels observed in testes and brain (Kan et al. 2005; Brawand et al. 2011; Soumillon et al. 2013; Mele et al. 2015). The high transcriptome complexity we observed in the pronephros may be a unique feature of this tissue and could indicate the presence of a more heterogeneous cell population or a more diverse set of isoforms among fewer cell types. More work is necessary to fully understand the function of the pronephros and why the transcriptome of this tissue is so diverse.

## Conclusions

By utilizing long-read RNA-sequencing, we were able to refine the gene annotations across five threespine stickleback tissues. Many of the Iso-Seq transcripts we identified were incorrect within the existing genome annotations or were completely novel. Furthermore, by using full-length transcripts, we identified extensive sex-specific and tissue-specific alternative splicing. Previous work with short-read sequencing has greatly underestimated the number of alternative isoforms. Overall, our results highlight the power of Iso-Seq for refining gene annotations and gaining a better understanding of transcriptome complexity across tissues.

## Methods

### Total RNA extraction, short-read, and long-read sequencing

All tissues were obtained from a laboratory-reared threespine stickleback fish, originally collected from the Japanese Pacific Ocean population (Akkeshi, Japan). Brain and liver samples were dissected from one adult male (one year old, 6.2 cm in standard length) and one adult female fish one year old, 6.3 cm in standard length). Pronephros samples were dissected from a separate adult male (one year old, 6.1 cm in standard length) and female fish (one year old, 6.1 cm in standard length). Gonads were dissected from a juvenile male (six months old, 4.6 cm in standard length) and a juvenile female (six months old, 4.8 cm in standard length). We selected juvenile stages to capture gonads that were actively undergoing meiosis (Craig-Bennett 1931; Borg and Van Veen 1982). Total RNA was extracted from all tissues using Trizol:chloroform RNA extraction, following the manufacturer recommended protocols (Invitrogen, USA). RNA from all tissues was used for both the Iso-Seq library preparation and the Illumina strand-specific RNA library preparation. Iso-Seq library preparation and sequencing was completed at the Georgia Genomics & Bioinformatics Core (University of Georgia, Athens, GA). All tissues were sequenced using a PacBio Sequel machine for 26 hours. Two SMRT cells were used for each tissue. Illumina strand-specific RNA library preparation and sequencing was completed by GENEWIZ (New Jersey, USA). Strand-specific libraries were sequenced on an Illumina HiSeq (2 × 150bp).

### Nuclei Isolation and ATAC-seq library preparation

Liver samples were collected from two juvenile males (~4.4 cm in standard length) and two juvenile females (~4.3 cm in standard length), originally collected from Lake Washington (Washington, USA). To isolate nuclei, half of the liver was homogenized in 1X PBS with proteinase inhibitor cocktail (PIC, cOmplete tablets Roche). The homogenized cells were then fixed using 16% formaldehyde and washed twice with PBS+PIC. The cells were lysed using a lysis buffer containing 1M Tris-HCl, pH=8, 0.5M EDTA, 10% NP-40, 50% glycerol/molecular grade H_2_O, and 1X PIC. Nuclei were counted using DAPI and then diluted to 60,000 – 80,000 nuclei. The ATAC-seq library preparation was performed using previously established protocols (Lu et al. 2017). To integrate the sequencing adapters, the diluted nuclei were rinsed with 1X TAPS to remove EDTA remaining from the lysis step. Tn5 transposase was then added to the nuclei and the reaction was carried out for 30 minutes at 37 °C. Immediately following the addition of sequencing adapters, the DNA was purified using a New England Biolabs Monarch DNA Cleanup kit (T1030). These library fragments were then amplified using Phusion enzyme (F530N) and Nextera PCR primers (Supplemental Table 7), using the following PCR conditions: 72 °C for 5 minutes, 98 °C for 2 minutes, and thermocycling at 98 °C for 10 seconds, 63 °C for 30 seconds, and 72 °C for 1 minute. Libraries were amplified for 13 cycles and Serapure beads were used to remove fragments smaller than 200 bp. ATAC-seq libraries were sequenced on Illumina NextSeq (2 × 150 bp) (Georgia Genomics & Bioinformatics Core, Athens, Georgia).

### Long-read RNA alignment and isoform identification

The eight tissues produced an average of 40.3 million raw subreads per tissue (583 gigabytes total; Supplemental Table 1). We analyzed the raw subreads following the Iso-Seq3 pipeline (v3.1; https://github.com/PacificBiosciences/IsoSeq). Circular consensus sequences (CCS) were created from the raw subreads using ccs (SMRTlink, v6; --noPolish --minPasses 1). We ran ccs on each tissue separately. cDNA primers (5’ AAGCAGTGGTATCAACGCAGAGTACATGGGG and 3’ AAGCAGTGGTATCAACGCAGAGTAC) were removed using lima (Iso-Seq3, v3.1; --isoseq --dump-clips --no-pbi). Nearly 70% of the CCS reads passed Lima default filters (Supplemental Tables 2 and 3). Trimmed circular consensus sequences were classified as full-length reads based on the presence of a poly-A tail using refine (Iso-Seq3, v3.1; --require-polyA). Full-length reads were clustered using cluster (Iso-Seq3, v3.1; default parameters). Any two full length reads were considered to be part of the same isoform if the 5` overhang was less than 100 bp long, the 3` overhang was less than 30 bp long, and any internal gaps between the reads were less than 10 bp long (https://github.com/PacificBiosciences/IsoSeq). These overlapping reads were clustered together. Clustered full-length reads were polished using polish (Iso-Seq3, v3.1; default parameters). The polished high-quality reads were aligned to the threespine stickleback genome (Ensembl build 97; (Jones et al. 2012b; Aken et al. 2016) using Minimap2 (v2.13) with the following parameters: -ax splice -uf –secondary=no -C5 (Li 2018). SAMtools was used to sort the alignments (Li et al. 2009; Li 2018). Reads that had accuracy scores less than 99% after polishing (low quality reads) were not considered further.

To remove redundancy among full length isoforms, identical isoforms were removed using cDNA Cupcake (collapse_isoforms_by_sam.py; https://github.com/Magdoll/cDNA_Cupcake). Count information for the reduced isoforms was then calculated using cDNA Cupcake (get_abudance_post_collapse.py, https://github.com/Magdoll/cDNA_Cupcake). An in-depth characterization of isoforms and removal of artifacts was completed using SQANTI (Tardaguila et al. 2018). SQANTI classified isoforms into nine different descriptors. Full splice matches (FSM) were defined as isoforms where all splice junctions fully matched an annotated gene. Incomplete splice matches (ISM) were defined as isoforms where some, but not all splice junctions matched. Isoforms were also categorized as genic intron (isoforms were located fully within an intron of an annotated gene), hereafter called intronic isoforms, genic genomic (isoforms overlapped an exon and an intron of an annotated gene) hereafter called genic, intergenic (isoforms were located outside of any annotated gene), fusion (isoforms spanned two annotated genes), and antisense (isoforms overlapped an annotated gene, but on the complementary strand). Novel isoforms of previously annotated genes were classified as either novel in catalog (NIC) or novel not in catalog (NNC). These categories were defined solely on whether the splice acceptors or donors for the splice junctions were known or novel. We combined isoforms from both the NIC and NNC categories in subsequent analyses. The default splice donors and acceptors in SQANTI were used (the most common sequences found in humans: GT-AG, GC-AG, and AT-AC; (Mount 1982; Ohshima and Gotoh 1987; Shapiro and Senapathy 1987; Tardaguila et al. 2018)). Stickleback-specific splice junction sequences are not known. Protein coding predictions were completed using GMST in SQANTI with the Ensembl transcriptome as the reference (Tang et al. 2015; Tardaguila et al. 2018). The SQANTI machine learning classifier identified artifacts based on features such as isoform length, presence of an open reading frame, the number of full-length reads per isoform, and any signature of reverse transcriptase switching (RTS), which results when the reverse transcriptase jumps across templates without stopping DNA synthesis. A true negative set was used to set the expectations for isoform artifacts. Because we did not have a true positive or negative set, we used the default behavior of SQUANTI, which assigns all the full splice match isoforms (FSM) as the true positives and the fusion isoforms as the true negatives. Isoform characterization was completed using sqanti_qc.py and filtering was completed using sqanti_filter.py. The filtered data set was rerun through sqanti_qc.py for the final characterization. The SQANTI filtered isoforms were used for the rest of the analyses.

### Assessing the completeness of each transcriptome

To assess whether our tissues were sequenced to an adequate depth, we utilized a subsampling approach (Workman et al. 2018). If our tissues were not sequenced sufficiently, the full set of isoforms may not have been captured. CCS reads were subsampled at 5%, 15%, 25%, 35%, 50%, 65%, 75%, 85%, and 95% of each individual tissue using picard (v2.16; DownsampleSam VALIDATION_STRINGENCY=SILENT) (http://broadinstitute.github.io/picard). The subsampled CCS reads were compared to the nucleotide sequences from the full tissue transcriptome using BLAST (v2.2.6, blastn, default parameters) (Altschul et al. 1990; Altschul et al. 1997; Camacho et al. 2009). The BLAST results for each tissue were filtered using custom python scripts. All BLAST alignments that covered at least 50% of the subsampled CCS read and at least 50% of an isoform from the full tissue transcriptome were retained. The total proportion of isoforms detected in each subsample compared to the full tissue transcriptome was calculated.

### Benchmarking universal single-copy orthologs (BUSCO)

To further assess transcriptome completeness, we utilized benchmarking universal single-copy orthologs (BUSCO, v3.0.2) (Simao et al. 2015; Seppey et al. 2019). BUSCO examines predicted genome annotations for completeness by using single-copy orthologs that are shared among metazoans. We used the metazoan (978 genes) and actinopterygii (4,584 genes) lineages for comparison (OrthoDB v9). The actionopterygii database was used because threespine stickleback fish are teleosts which is the largest infraclass of actinopterygii. The predicted amino acid sequences for all protein coding genes for the full transcriptome after SQANTI filtering were inspected (BUSCO gene set assessment). Protein coding genes from build 97 from Ensembl were also assessed. The zebrafish *(Danio rerio)* was set as the default species and all other parameters were left at the default settings.

### Comparisons between transcriptomes

For analyses, we assembled several different combinations of transcriptomes using the dataset create function of SMRTlink (v. 6). To examine the differences between the Ensembl transcriptome and the transcriptome produced from the Iso-Seq data, we utilized all eight tissues (hereafter referred to as the Iso-Seq transcriptome). To examine if any isoforms were sex-specific, all five tissues (brain, liver, pronephros, testis, and ovary) were combined for each sex (hereafter referred to as the female transcriptome and the male transcriptome). We also examined sex-specificity in somatic tissues only, combining only the brain, liver, and pronephros of each sex (hereafter referred to as the somatic female transcriptome and the somatic male transcriptome). Lastly, we compared transcriptomes of individual tissues. All combined transcriptomes and the individual tissue transcriptomes were subject to the same Iso-Seq3 pipeline beginning with the removal of cDNA primers with Lima.

We assigned universal isoform identifications for the full Iso-Seq transcriptome that combined all tissues. BLAST was used to compare isoforms among individual tissue transcriptomes (v2.2.6, blastn, default parameters) (Altschul et al. 1990; Altschul et al. 1997; Camacho et al. 2009). Duplicate isoforms within the full Iso-Seq transcriptome were first removed by identifying any isoforms that aligned exactly to another isoform (i.e. BLAST alignments were identical between the isoforms). This removed 1,139 isoforms from the full Iso-Seq transcriptome. All other isoforms in each transcriptome (i.e. individual tissues, female transcriptome, male transcriptome, somatic female transcriptome, or somatic male transcriptome) were compared to this full transcriptome with BLAST. The BLAST results were filtered using custom python scripts. A positive alignment was identified if at least 50% of the query sequence matched at least 60% of the subject sequence. Query isoforms that matched more than one subject isoform were collapsed to a single isoform, keeping the longest alignment. Any isoforms that did not meet these criteria were discarded.

### Short read RNA alignments

For short read RNA sequencing, low quality regions and residual adapters were removed using trimmomatic (v0.36) with default parameters including ILLUMINACLIP (Bolger et al. 2014). Trimmed reads were aligned to the threespine stickleback genome (Ensembl build 97 (Jones et al. 2012b; Aken et al. 2016) using Hisat2 (v2.1) with the following parameters: phred33, rna-strandness FR (Kim et al. 2015). Short reads were utilized by SQANTI to examine the overall expression level of individual isoforms and to verify splice junctions (Tardaguila et al. 2018). Gene expression matrices with TPM (transcripts per million) were calculated using Kallisto (v0.46) (Bray et al. 2016). The quant function of Kallisto was used on trimmed short reads from each tissue and the combined data sets. To calculate the read coverage across splice junctions, STAR (v2.7, default parameters) was run on each tissue (Dobin et al. 2013). The output from Kallisto and STAR were then used an input for the sqanti_qc.py.

### ATAC-seq genome coverage at TSSs

Residual adapter sequences from the Nextera primers were trimmed using trimmomatic (v0.36) with PE and ILLUMINACLIP (keeping both reads) and the following parameters: LEADING:0 TRAILING:0 MINLEN:30 (Bolger et al. 2014). Trimmed reads were aligned to the revised threespine stickleback genome (Nath et al. 2020), including the mitochondrion and unplaced scaffolds, using Bowtie2 (v2.3.5; -X 1000 –no-unal) (Langmead and Salzberg 2012). Reads with a mapping quality less than 20 were filtered from the alignments using SAMtools (v1.10) (Li et al. 2009). PCR duplicates were removed using MarkDuplicates from Picard (v2.21.6, default parameters). The read coverage per bp was calculated using Bedtools (v2.26, genomecov -d) (Quinlan and Hall 2010). Custom python scripts were used to average the read coverage across a 4kb window surrounding the Ensembl and Iso-Seq TSSs.

### Characterizing Non-coding RNAs by size and genome location

Non-coding RNAs (ncRNAs) were characterized by size and genome location using custom python scripts. All ncRNAs were defined as isoforms that did not have detectable protein coding potential, using the GeneMarkS-T algorithm within SQANTI (Tang et al. 2015; Tardaguila et al. 2018). ncRNAs are generally classified based on overall length: short ncRNAs are less than 200 bp and long ncRNAs are greater than 200bp (Jacquier 2009; Pauli et al. 2011). We separated ncRNAs into these two length categories as well as three main classes: intergenic, intronic, or antisense (Pauli et al. 2012). Intergenic ncRNAs were ncRNAs that did not overlap with any genes. Intronic ncRNAs were ncRNAs that fell completely within an intron of another gene and shared no overlap with the surrounding exons. Antisense ncRNAs were ncRNAs that did overlap with a gene but were on the opposite strand. Any ncRNA that was not classified into one of these three categories was added to an unknown category. To characterize all ncRNAs into one of these three classes, we compared the location of each ncRNA to all annotated Ensembl transcripts and the newly identified Iso-Seq isoforms.

### Novel genes protein domain search through InterProScan

We used InterProScan (v.5.32) (Jones et al. 2014) to identify protein domains and potential functions of novel proteins. The amino acid sequences from all novel protein coding genes from the full Iso-Seq transcriptome were used. All available databases in IntroProScan were used to compare protein sequences. InterProScan was run with default parameters and GO terms and pathway information was recorded.

### Gene Ontology Analysis

Gene ontology (GO) terms enrichment analysis was completed using custom python scripts. Gene IDs and GO terms were downloaded from Ensembl using Biomart (Smedley et al. 2015). GO terms of the novel genes that were identified through InterProScan were also added to the complete list of GO terms for threespine stickleback. The total number of occurrences for each GO term was calculated for several analyses. GO term enrichment in each set was compared against 10,000 random permutations of the same sample size randomly drawn from the total set of genes. P-values were adjusted for multiple testing using a Bonferroni correction based on the total number of observed GO terms in each set. Enriched GO terms were visualized using web gene ontology annotation plot (WEGO) (Ye et al. 2006; Ye et al. 2018).

### Data Access

The project has been deposited at the NCBI Sequence Read Archive under two BioProjects. All raw Iso-Seq subread bam files and short read RNA-seq fastq files are deposited in BioProject PRJNA633846. The liver ATAC-seq fastq files are present in BioProject PRJNA667175.

## Supporting information

Supplemental File 1: Classification file

Supplemental File 6: Sex-specific transcripts

Supplemental File 7: Tissue-specific transcripts

## Acknowledgements

This research was funded by the National Science Foundation IOS 1645170 (M.A.W.), the National Science Foundation MCB 1943283 (M.A.W.), the Office of the Vice President of Research at the University of Georgia (M.A.W.), the Jan and Kirby Alton Fellowship (Department of Genetics, UGA to A.S.N), and the Rosemary Grant Award (Society for the Study of Evolution to A.S.N). We also thank Robert Schmitz and his lab at UGA for assistance and reagents for the ATAC-seq protocol. PacBio Sequencing and library preparations were conducted at the Georgia Genomics and Bioinformatics Core at the University of Georgia. All custom scripts are publicly available on Github under ASNaftaly: https://github.com/ASNaftaly/IsoSeq3_Stickleback.

