## Supplemental Figures for "Long-read RNA sequencing reveals widespread sex-specific alternative splicing in threespine stickleback fish"

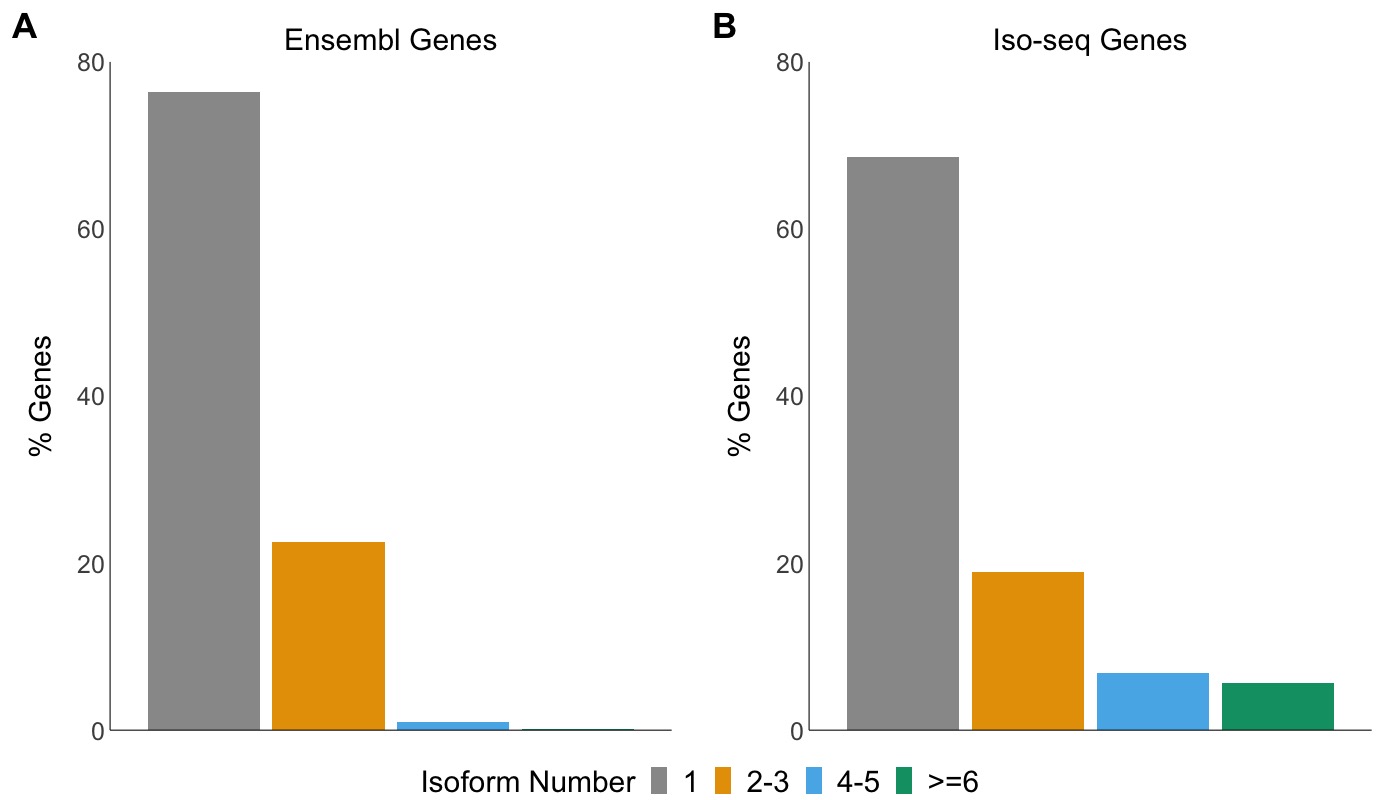

Supplemental Figure 1. There are more isoforms per gene in the Iso-Seq transcriptome compared to the Ensembl transcriptome. (A) In the Ensembl transcriptome, only 24% of genes have more than one isoform. (B) In the Iso-Seq transcriptome, 31% of all genes have at least two isoforms.

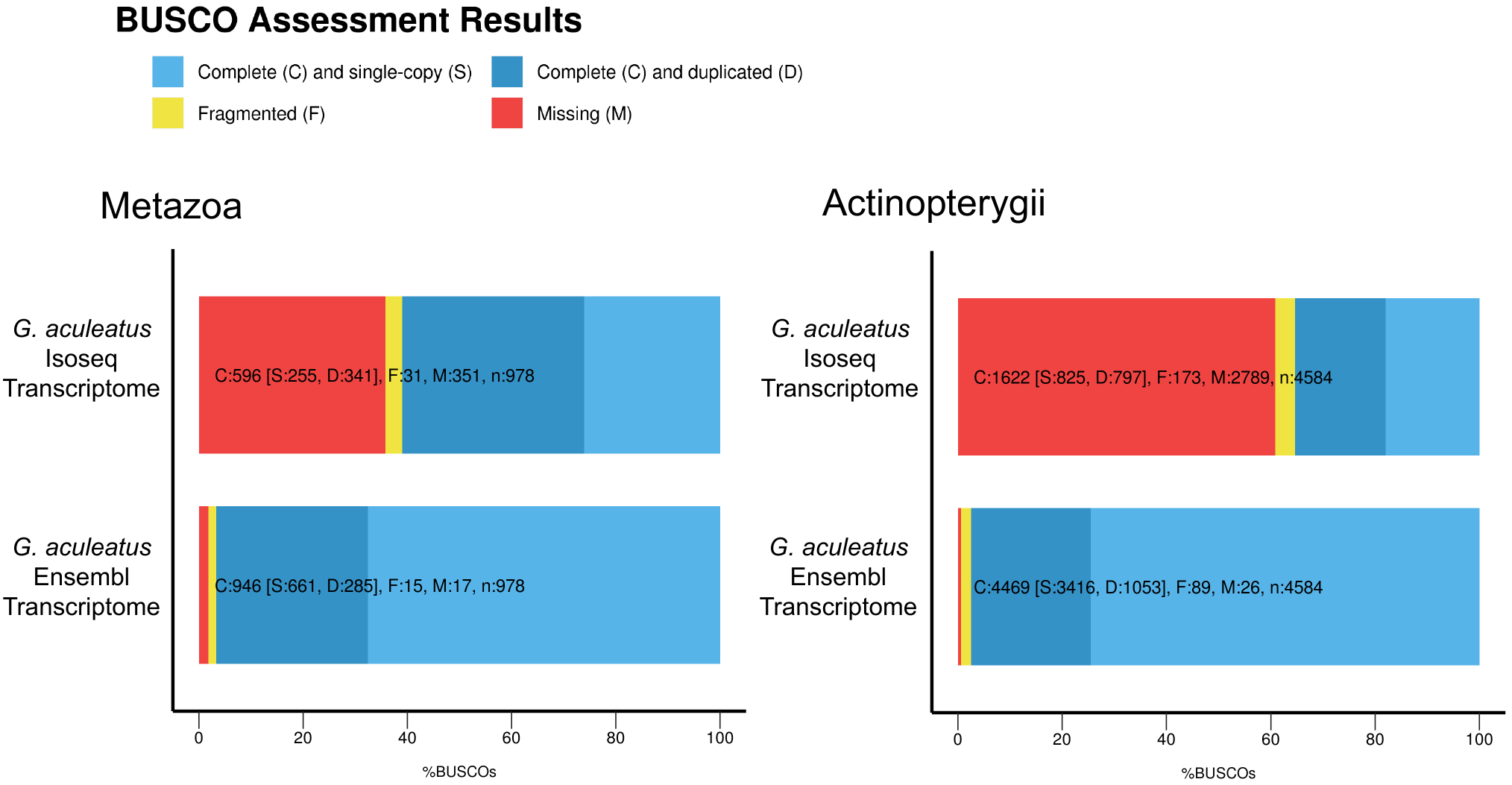

Supplemental Figure 2. The Iso-Seq transcriptome contained fewer complete BUSCO orthologs compared to the Ensembl annotations. The Ensembl transcriptome is almost complete in Metazoan (97% complete orthologs) and Actinopterygian (98% complete orthologs) lineages. The Iso-Seq transcriptome, while not complete, represents 61% of complete Metazoan orthologs and 35% of complete Actinopterygian orthologs.

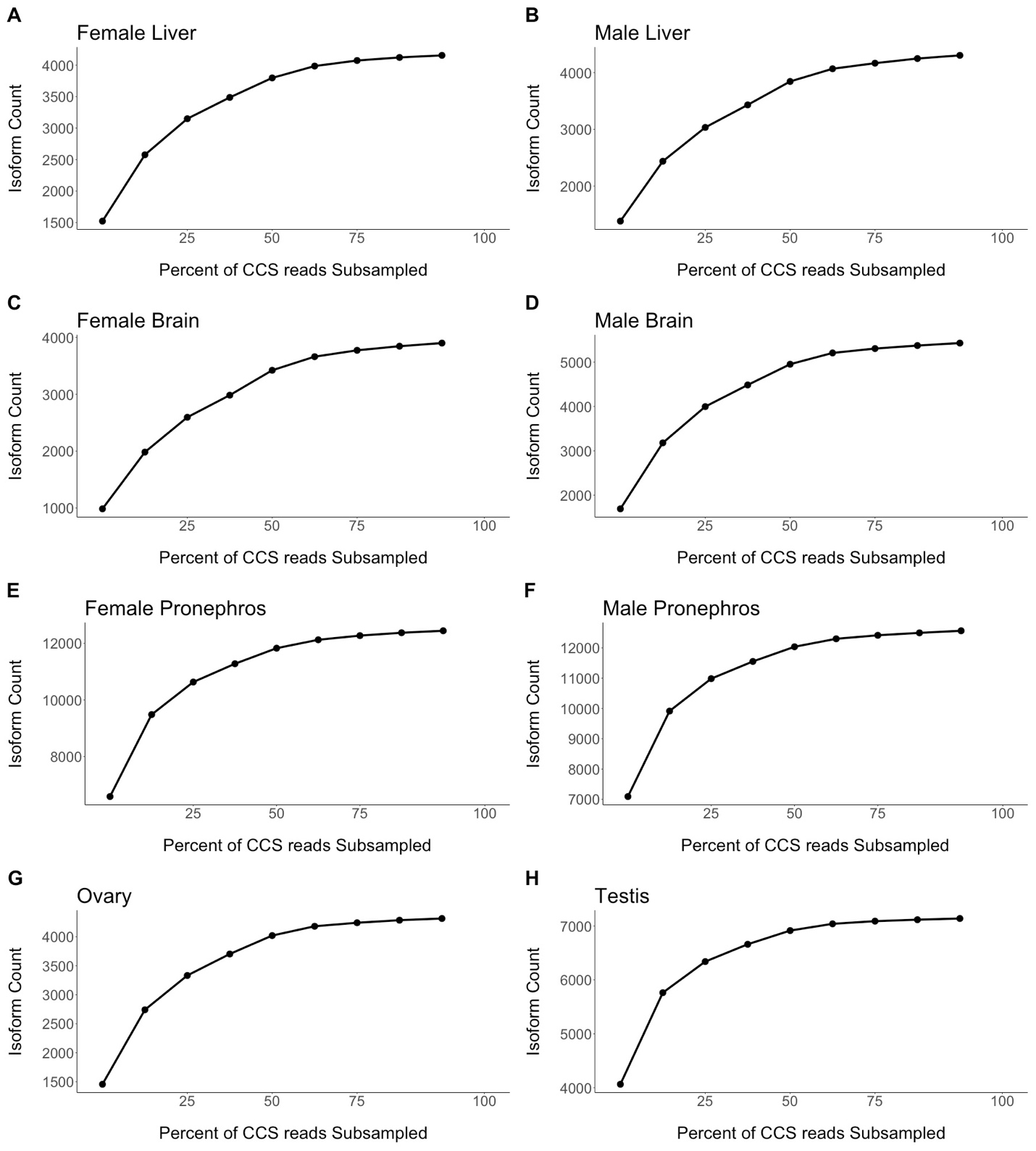

Supplemental Figure 3. Over 90% of total isoforms are recovered when subsampling CCS reads. 90% of total isoforms are recovered at the following percent of subsampled CCS reads: (A) 65% for the female liver, (B) 85% for the male liver, (C) 85% for the female brain, (D) 65% for the male brain, (E) 65% for the female pronephros, (F) 50% for the male pronephros, (G) 65% for the ovary, and (H) 35% for the testis.

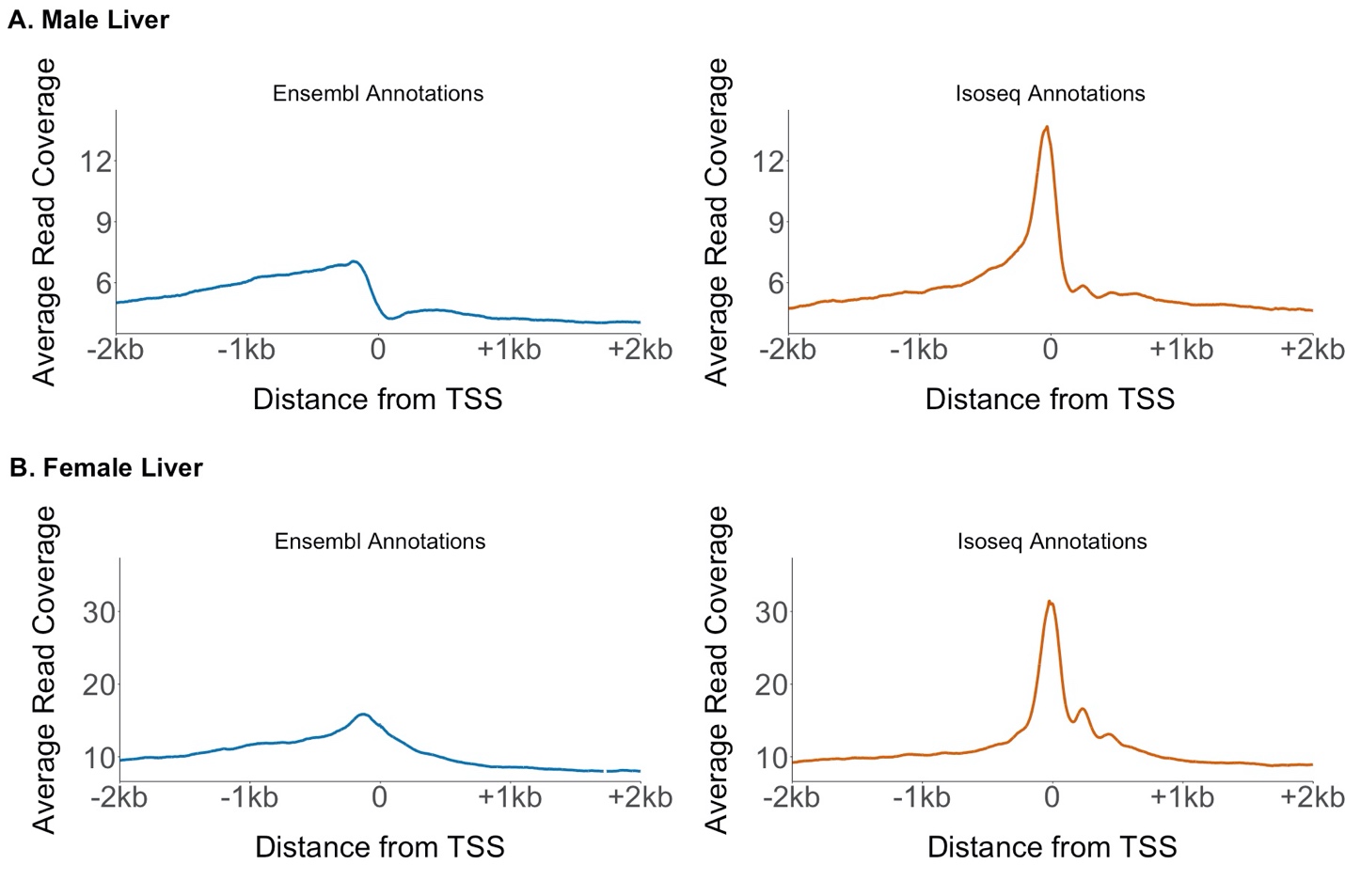

Supplemental Figure 4. Accessible chromatin is localized in narrow peaks around the Iso-Seq transcription start sites. We compared ATAC-seq read coverage at all Ensembl TSSs and Iso-Seq TSSs across the autosomes. ATAC-seq reads show an enrichment at the Iso-Seq TSS compared to the Ensembl TSS. This indicates a more accurate positioning of the TSS using Iso-Seq. A second male and female replicate is shown in Figure 3.

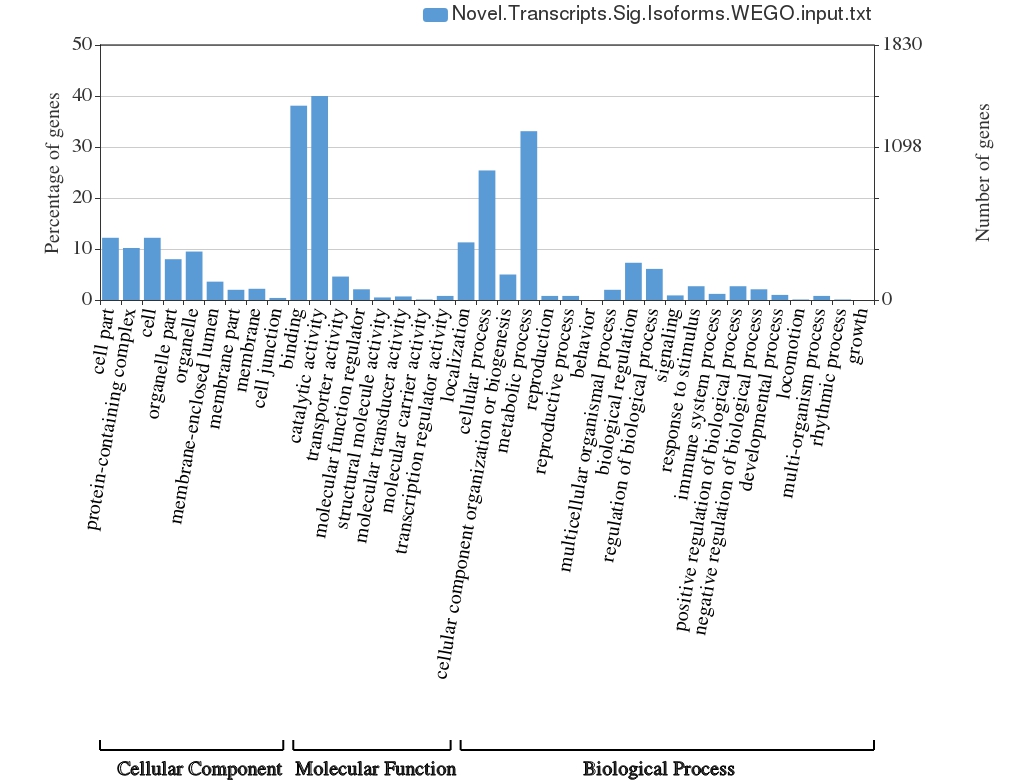

Supplemental Figure 5. Novel isoforms have many general cellular functions. 595 GO terms were significantly enriched (P < 0.001) after a Bonferroni correction. GO terms were collapsed into more general categories using WEGO (web gene ontology annotation plot).

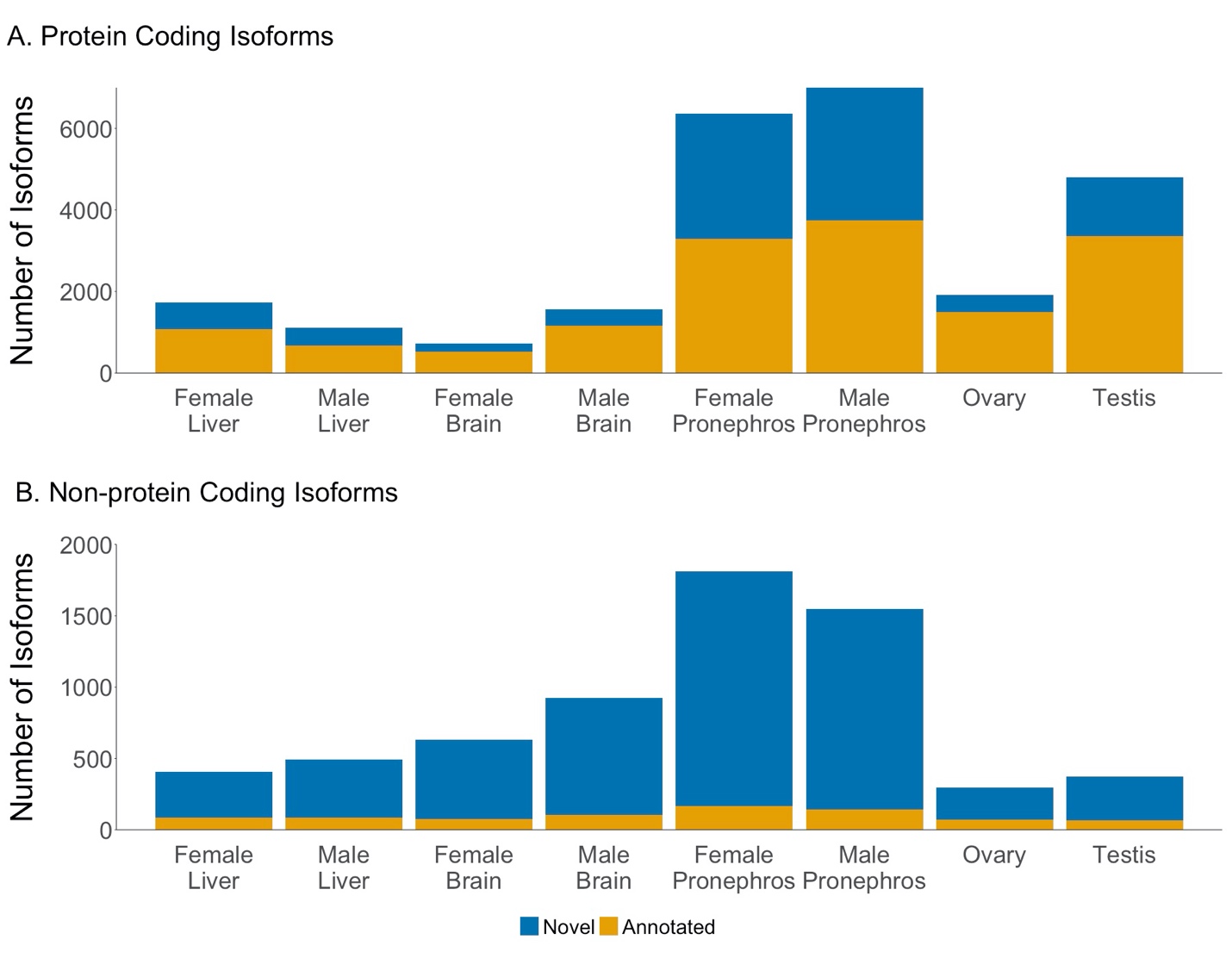

Supplemental Figure 6. Most of the novel isoforms are non-protein coding. (A) Most protein-coding isoforms were previously annotated in the Ensembl transcriptome. The pronephros and the testis have the largest number of novel isoforms. (B) A majority of non-protein coding isoforms across all samples are novel isoforms.
